# Soil composition and plant genotype determine benzoxazinoid-mediated plant-soil feedbacks in cereals

**DOI:** 10.1101/2021.04.14.439871

**Authors:** Selma Cadot, Valentin Gfeller, Lingfei Hu, Nikhil Singh, Andrea Sánchez-Vallet, Gaétan Glauser, Daniel Croll, Matthias Erb, Marcel G. A. van der Heijden, Klaus Schlaeppi

## Abstract

Plant-soil feedbacks refer to effects on plants that are mediated by soil modifications caused by the previous plant generation. Maize conditions the surrounding soil by secretion of root exudates including benzoxazinoids (BXs), a class of bioactive secondary metabolites. Previous work found that a BX- conditioned soil microbiota enhances insect resistance while reducing biomass in the next generation of maize plants. Whether these BX-mediated and microbially driven feedbacks are conserved across different soils and response species is unknown. We found the BX-feedbacks on maize growth and insect resistance conserved between two arable soils, but absent in a more fertile grassland soil, suggesting a soil-type dependence of BX feedbacks. We demonstrated that wheat also responded to BX-feedbacks. While the negative growth response to BX-conditioning was conserved in both cereals, insect resistance showed opposite patterns, with an increase in maize and a decrease in wheat. Wheat pathogen resistance was not affected. Finally and consistent with maize, we found the BX-feedbacks to be cultivar specific. Taken together, BX- feedbacks affected cereal growth and resistance in a soil and genotype dependent manner. Cultivar-specificity of BX-feedbacks is a key finding, as it hides the potential to optimize crops that avoid negative plant-soil feedbacks in rotations.

## Introduction

Plants modify their surrounding soil environment to optimize their performance and these soil modifications then affect the performance of other plants growing later in this soil. This phenomenon, where soil legacies from a previous plant generation modify the performance of the next plant generation, are well-known as plant-soil feedbacks (PSFs, van der Putten et al., 2013). The first plant generation may change physical, chemical and/or biological characteristics of the soil, a process often referred to as “soil conditioning” (Bezemer et al., 2006). Such legated modifications of the soil condition then feedback on the next generation’s growth, development and/or metabolism, and can influence defence and/or tolerance to biotic and abiotic stresses (Huberty et al., 2020; Pineda et al., 2020; Revillini et al., 2016; Zhao et al., 2021).

Negative PSFs are thought to occur because the previous plant generation enriched pathogens, released allelopathic compounds and/or reduced the availability of nutrients whereas a promotion of beneficial symbionts and/or an enhancement of nutrient availability by the previous generation would result in positive PSFs (Bever, 2003; van der Putten et al., 2013). Therefore, PSFs represent important factors that determine the coexistence and diversity of plant communities in natural ecosystems (Bennett et al., 2017; Putten, 2017; Teste et al., 2017). For instance, PSFs regulate through mycorrhizal symbiont types the population structure of temperate forests (Bennett et al., 2017). Interestingly, not only the conditioning by the first plant generation, but also the ’interpretation of soil legacies’ by the subsequent plant generation is key that PSFs promote plant diversity. Teste et al. (2017) revealed in Mediterranean shrublands that the nutrient-acquisition strategy of the response plants explained differential feedbacks to soil biota and thereby promoted local plant diversity. Besides in natural ecosystems, PSFs are very important in agricultural context where the growth of previous crop impacts the performance and yield of the following crop (Mariotte et al., 2018; Pizano et al., 2019). Although, not specifically following the terminology of PSFs, farmers have recognized since a long time that certain sequences of crop plantings negatively impacted their yields. Using dedicated crop rotations, farmers take advantage of positive while avoiding negative PSFs. For example, *Brassica* crops are specifically used in rotations, a cropping method referred to as biofumigation, because of their positive feedbacks on the following crops (Gimsing and Kirkegaard, 2009). *Brassicaceae* plants produce secondary metabolites, i.e. glucosinolates, that function as natural biocides to control fungal and oomycete pathogens and plant-parasitic nematodes (Brennan et al., 2020; Poveda et al., 2020). Coupling such practical knowledge with a fundamental ecological understanding is likely to offer further opportunities for harnessing PSFs in sustainable agriculture (Mariotte et al., 2018; Pineda et al., 2020). Possible applications of PSFs ranging from seed selection to agricultural biodiversity management, including fine-tuning the already widely used crop rotation systems.

PSFs often affect the next plant generation’s defence against pests and pathogens. For example, soil conditioning by eight forb and grass species consistently suppressed aboveground infestations by *Frankliniella occidentalis* where these plants shaped species-specific soil microbiomes causing varying amplitudes of thrips resistance, secondary metabolite concentrations in leaves and plant growth (Pineda et al., 2019). These positive PSFs on resistance against thrips were not effective against the spider mites *Tetranychus urticae*, corroborating that differing plant traits shape specific soil legacies that then result into specific plant–insect feedbacks (Heinen et al., 2020). Similarly, PSFs improved plant protection against pathogens as for example in *Arabidopsis thaliana* against the aboveground pathogen *Pseudomonas syringae* (Yuan et al., 2018) or against the root pathogen *Pythium ultimum* shown with Chrysanthemum by (Hannula et al., 2020). Pest and pathogen control in PSFs may function via attraction or enrichment of beneficial soil microbes or their shifts in the whole microbial community and they function in positive feedbacks (Bakker et al., 2018; Berendsen et al., 2018).

Plants secrete considerable amounts of bioactive molecules from their roots into the rhizosphere and this root exudation is responsible for a large part for belowground soil conditioning (Bever et al., 2012). Exudation directly benefits a plant by enhancing nutrient availability (Jones and Darrah, 1995), suppressing competitors by allelopathy (Inderjit and Duke, 2003) and importantly, it is largely responsible for the plant’s influence on the rhizosphere microbiota, which then indirectly benefits the plant (Sasse et al., 2018). The rhizosphere microbiome confers health benefits to the plant (Berendsen et al., 2012), by mechanisms such as direct protection (antibiosis), niche competition with pathogens for resources or by enhancing the immune response from the plant via induced systemic resistance (ISR, Pieterse et al., 2014). Root exudate metabolites such as coumarins or benzoxazinoids are known to selectively structure the rhizosphere microbiome, and these changes have been linked to improved plant health (Hu et al., 2018; Stringlis et al., 2018).

Benzoxazinoids (BXs) are a family of secondary metabolites, produced by plants from the Poaceae family, including important cereals such as wheat, maize and rye. The main BX compound exuded by maize and wheat is the glucoside 2,4- dihydroxy-7-methoxy-2H-1,4-benzoxazin-3(4H)-one (DIMBOA-Glc), which is deglycosylated and degraded to 6-methoxy-benzoxazolin-2-one (MBOA; Niculaes et al., 2018) in soil. MBOA is a bioactive catabolite with allelopathic properties (Macías et al., 2009; Schandry and Becker, 2020), and has also direct protective functions against herbivore insects (Niemeyer, 2009; Wouters et al., 2016; Zhou et al., 2018) and fungal or bacterial pathogens by toxicity (Martyniuk et al., 2006; Schalchli et al., 2012), or by suppressing the virulence of pathogens (Sicker et al., 2000). In soil, MBOA is further degraded by microorganisms to the more stable and highly allelopathic 2-Amino-7-methoxy-3H-phenoxazin- 3-one (AMPO) (Etzerodt et al., 2006; Macías et al., 2009; Schandry and Becker, 2020). BXs function in selectively structuring the root and rhizosphere microbiomes (Cadot et al., 2020; Cotton et al., 2019; Hu et al., 2018; Kudjordjie et al., 2019), as shown by experiments comparing BX-producing wild type and BX-defective mutant plants. The selective structuring of microbial communities functions not only via antagonistic functions of BXs described above, but also through attraction, as evidenced by the positive chemotactic response of *Pseudomonas putida* to locate maize roots (Neal et al., 2012). Of note, the selective recruitment of *P. putida* by BXs may promote plant health, as this strain is capable of triggering ISR in maize (Neal and Ton, 2013).

The secretion of BXs conditions the surrounding rhizosphere microbiota that then drives PSFs on the next plant generation (Hu et al., 2018). These feedbacks become visible when comparing the soil variant that was conditioned by BXs (**BX+ soil**) with the control soil variant that was not conditioned by BXs (**BX- soil**). To condition the soil, wild-type and BX exuding maize plants such as the inbred line B73 are grown in the first generation. The control soil is prepared by cultivating alongside BX mutants such as the near isogenic line *bx1*(B73), which is compromised in the biosynthesis and secretion of BXs (Maag et al., 2014). Feedbacks on a new generation of maize plants included enhanced insect resistance and reduced plant growth for plants grown on BX+ soil. The maize plants responded physiologically with an increase in defence hormone levels and an upregulation of defence marker genes. Sterilization and complementation experiments demonstrated that these feedbacks were driven by the soil microbiota. We refer to these BX-mediated and microbially driven PSFs hereafter as ‘BX-feedbacks’ for simplification.

A number of follow-up questions emerged from our initial mechanistic study on BX-feedbacks, which was limited to maize and examined only a single soil, the clay loam ‘Changins soil’. It is well-known that different soil types and microbiota pools influence PSFs (Bergmann et al., 2016; Smith-Ramesh and Reynolds, 2017), and we also know that BX-sensitive microbes differ between different soils (Cadot et al., 2020). Therefore, we asked whether BX-feedbacks would also occur in other soils. It is also common that PSF responses vary between species and cultivars (Heinen et al., 2020; Kuťáková et al., 2018; Wagg et al., 2015), including different maize lines (Hu et al., 2018). However, it is unknown how another plant species than maize would respond to a BX-conditioned soil. Wheat is an important crop that is often following maize in crop rotations in Europe and therefore, its BX-feedbacks are particularly relevant from an agricultural point of view. The third emerging question was related to the defensive range of BX- feedbacks, as it was tested only on insect resistance so far. Because of reported trade-offs between pathogen and insect defence strategies (Erb et al., 2011; Zhu et al., 2018), we asked whether BX-feedbacks would also impact pathogen defence. Finally, we tested whether BX-feedbacks could be preserved by storing conditioned soils in the cold room, as a simulation of overwintering. In this study, we addressed the above mentioned open questions and we showed that BX-feedbacks are also effective on wheat and that they function in a soil and cultivar-dependent manner.

## Methods

### Overview of BX-feedback experiments

This study comprised six complementary experiments, all investigating BX- feedbacks comparing BX-conditioned (**BX+ soil**) with non-conditioned soil variants (**BX- soils**; see below). ‘Experiment 1’ was designed to study BX- feedbacks in another soil and measured BX-feedbacks on growth and insect defence of wild-type maize in ‘Reckenholz’ soil, along with ‘Changins’ soil as a control because BX-feedbacks were originally discovered in the latter (Hu et al., 2018). With the ‘Reckenholz’ soil we further compared freshly conditioned soil with soil that was conditioned a year before. Goal was to learn, whether BX- feedbacks remain preserved upon storage of the conditioned soils for up to one year at 4°C in the cold room. Similarly, we tested the additional soil ‘Q-matte’ relative to ‘Changins’ soil for BX-feedbacks on maize in ‘Experiment 2’. The ‘Experiments 3 to 5’ were setup to investigate if BX-feedbacks also occur on wheat. We performed these experiments mainly with ‘Changins’ soil because the feedbacks on maize were originally discovered with this soil (Hu et al., 2018). In experiment 3’ we examined BX-feedbacks on growth (biomass) and physiology (chlorophyll content, hormones) testing two wheat cultivars as response plants. For comparison, we included Reckenholz soils batches as used in the first experiment. ‘Experiment 4’ and **‘**Experiment 5’ were designed to specifically test BX-feedbacks on insect and pathogen resistance, respectively. We conducted performance assays with the herbivore *Spodoptera frugiperda* with plants of two wheat cultivars in ‘Experiment 3’. With ‘Experiment 5’ we studied resistance against the wheat pathogen *Zymoseptoria tritici* using one of the two cultivars. Finally, ‘Experiment 6’ was conducted with potting soil to uncover if the genotype-dependent BX-feedbacks on maize shoot biomass and insect resistance occur during the conditioning or the feedback phase. See the **Supplementary Information** for the detailed experimental setups of each of these six experiments.

### Feedback experiments with conditioned soils

BX-feedback experiments rely on the comparison of soil variants that were *conditioned by BXs* (**BX+ soil**) with variants that were *not conditioned by BXs* (**BX- soil**) as control. Conditioning functioned by growing the wild-type (WT) line B73, which secreted BXs to the surrounding soil (Hu et al., 2018; Maag et al., 2014). The control soil variant was prepared by growing the near isogenic mutant line *bx1*(B73), which is compromised in the biosynthesis and secretion of BXs (Maag et al., 2014). Soil variants conditioned by growth of WT maize are referred to as **BX+ soil**, and soil variants, where the mutant *bx1* was grown, are referred to as **BX- soil**. Soil conditioning was standardised, by growing the B73 and *bx1* lines for 3 months in a complete randomized block design.

For this study we used several batches of conditioned soils and they were either prepared in field or in greenhouse experiments (**Table S1**, detailed below). We have previously described the details of the field experiments, of which we collected the conditioned soil batches ‘Changins 2016’ (Hu et al., 2018) and ‘Reckenholz 2016’ (Cadot et al., 2020). The conditioned soil batch ‘Reckenholz 2015’ (**Table S1**) was harvested from a field experiment, which we performed in 2015 with the same setup as the field experiment in 2016. See the **Supplementary Information** for the details of the field experiment conducted at Agroscope Reckenholz in 2015. We needed to condition additional soils for further experiments (**Table S1**) and this was done in the greenhouse, knowing that this type of conditioning works, too (Hu et al., 2018). The details of the greenhouse experiments to condition the soil are documented in the Supplementary Information.

Prior to their use for feedback experiments, all batches of conditioned soils were sieved to 1 cm and at the same time mixed with autoclaved quartz sand in a 1:4 sand to soil proportion (20% by volume). Importantly, the metal sieve was always sterilized with 70% ETOH between BX+ and BX- soil variants to avoid transfer of soil and microbes.

We have previously reported the chemical and physical characteristics from the clay loam soil in Changins and loam soil from Reckenholz (Cadot et al. 2020). We also determined the soil characteristics of the silt loam soil from Q-Matte in the same laboratory using the same certified methods (Labor für Boden- und Umweltanalytik, Eric Schweizer AG, Thun, Switzerland). Soil texture classes were determined using the online tool of the Natural Resources Conservation Service for Soils at the Unites States of Department of Agriculture. See **Table S2**.

### Chlorophyll measurements

Wheat chlorophyll content was measured in Experiments 3 and 4 on the second fully opened leaf using a Soil Plant Analysis Development SPAD-502 meter (Minolta Camera Co., Japan), at around 1 cm below the tip on two plants per pot, and the values were averaged for each pot for analysis.

### Phytohormone analysis

To obtain insights into the plant defence status, we measured the following plant hormones (or precursors) in wheat samples of Experiment 3 salicylic acid (SA), oxophytodienoic acid (OPDA), jasmonic acid (JA), jasmonic acid-isoleucine (JA-Ile) and abscicic acid (ABA). The frozen leaf material was ground using mortar, pestle and liquid nitrogen, transferred to 1.5 mL microcentrifuge tubes and precisely weighed. Hormone analysis was performed according to (Glauser et al., 2014). Resulting hormone concentrations were then standardized according to initial leaf weight.

### Insect assays

Second (Experiment 6) or third (Experiment 1) instar *Spodoptera frugiperda* caterpillars were used to assess maize insect resistance as described in Hu et al. (2018). Briefly, they were weighed and selected for similar weights before starting the feeding assay. They were placed into transparent plastic cups (4 cm height and 3.5 cm diameter) that have perforated plastic lids. The youngest fully developed leaves of individual maize plants were used for feeding. Leaves were placed as ca. 3 cm portions (excluding the leaf tip) into the cups with some moisture. These leave segments were daily replaced by new portions of the same leave throughout the assay. Caterpillar mass was determined 3 days after the start of the assay and the growth rate calculated (Growth rate = (Weight at day X – initial weight) / initial weight).

In Experiment 3 we evaluated wheat insect resistance using 2^nd^ instar caterpillars of *Spodoptera littoralis*. They were weighed before starting the feeding assay and two to three caterpillars were placed in small perforated plastic boxes. Two leaves per wheat pot were fed portioned by pieces for one week. Leaf pieces were replaced in the box and moisturized after three days and then every two days as the caterpillars grew bigger. Caterpillars were weighed at 4, 5 and 7 days (D) of feeding and the growth rate was calculated (formula as above).

### Pathogen assay

In experiment 4, we tested for eventual differences in pathogen resistance of the wheat cultivar Drifter when growing on ‘BX+’ or ‘BX- soils’ using the fungus *Z. tritici* (strain 3D7). The second fully unrolled leaf of 3-week-old plants were spray-infected with 10 ml of blastospores following a standard infection protocol (Singh et al., 2021). The inoculated plants were kept at 100% relative humidity and 21^0^C for 2 days, before going back to initial growth conditions (see above). Five additional plants were left uninoculated as controls. The inoculated leaves were harvested 21 days post infection for counting pycnidia (reproductive structures of the fungus) using the software ImageJ.

Fungal biomass in the inoculated leaves was quantified on the same plant material by quantitative PCR (qPCR), using the fungal strain-specific forward primer sequence 5’cgacatcggttcagagatggaa‘3 and reverse primer sequence 5’gtaccttcgattcgtgcggt‘3, and the plant 18S forward primer sequence 5’cgcagcaaatcccacgg’3 and reverse primer sequence 5’gcgcagcttcttccactttgac’3. Genomic DNA was extracted with a DNA easy plant kit (Qiagen, Germany) with minor modifications (samples were ground on the grinding machine Qiagen Tissue Lyser II, 2 x 30s at setting 30, incubated in the final step for 15 min at 65°C and the DNA was eluted in a volume of 150 ul). Real time qPCR reactions were then performed according to Meile et al. (2018) and Barrett et al. (2021). We used the ΔΔC_T_ method to express the fungal biomass relative to the plant leaf signal (E^ΔC_T_ (Fungus, control – infected)/ E^ΔC_T_ (Plant, control – infected).

### Statistical analyses

Data analysis was performed in R (version 3.5.1, R Development Core Team, 2017) using the package ‘ggplot2’ for plotting (Wickham, 2016). All statistical code (incl. the models used) and the design of Experiments 1 to 6 are documented in **Data S1** to **S6**, respectively. As general approach, we inspected the data whether they satisfied normality assumptions using residual plots following (Fahrmeir et al., 2013). Most one-time point data (biomass, height, chlorophyll, hormones) was examined with ANalyses Of VAriance assessing the effect of *conditioning* (BX+ vs. BX-) and combinations with effects of *soil* (Experiment 1 & 2), *wheat line* (Experiment 3 & 4), or *genetic background* (Experiment 6). Pair-wise T-tests were performed in a post-hoc manner (Experiment 1, 2, 3 and 4) or using Tukey HSD tests (Experiment 6). Data with multiple time points (4x SPAD in Experiment 4; 3x caterpillar in Experiment 5) were analysed with linear mixed-effect models (LMM) using the package ‘lme4’ (Bates et al., 2015). Pycnidia counts from Experiment 5 were analysed with generalized linear model (glm) using a quasipoisson distribution model.

## Results

### BX-feedbacks on maize are soil-dependent

To answer the first major question of this study - Do BX-feedbacks on maize also exist in other soils? - we tested with two experiments whether the BX- feedbacks, as observed in Changins soil (Hu et al., 2018), also occur in two other soils. Reckenholz and Q-Matte are loam and silt loam soils, respectively and have markedly different physicochemical characteristics compared to clay loam soil in Changins (**Table S2**). We grew B73 and *bx1*(B73) plants to condition the ‘BX+’ and ‘BX-‘ soil variants for subsequent feedback experiments, where we then measured shoot height, shoot biomass and insect resistance in the next generation of maize plants. In the first experiment we compared Reckenholz relative to Changins soil. Consistent with the Changins soil, we also found for the Reckenholz soil a significantly lower shoot biomass (**Fig. 1A**) and shoot height (**Fig. 1B**) when plants were grown on ‘BX+’ compared to ‘BX-‘ soil variants (**Data S1**, documents the statistical analyses). Biomass reduction on ‘BX+ soil’ accounted for -43.7 % in the Reckenholz soil, which is similar as for the Changins soil (-46.1%). Upon testing insect resistance, we found that *S. frugiperda* caterpillars grew at a reduced rates in Reckenholz (-32.2 %) and Changins (-29.6

**Figure 1.**
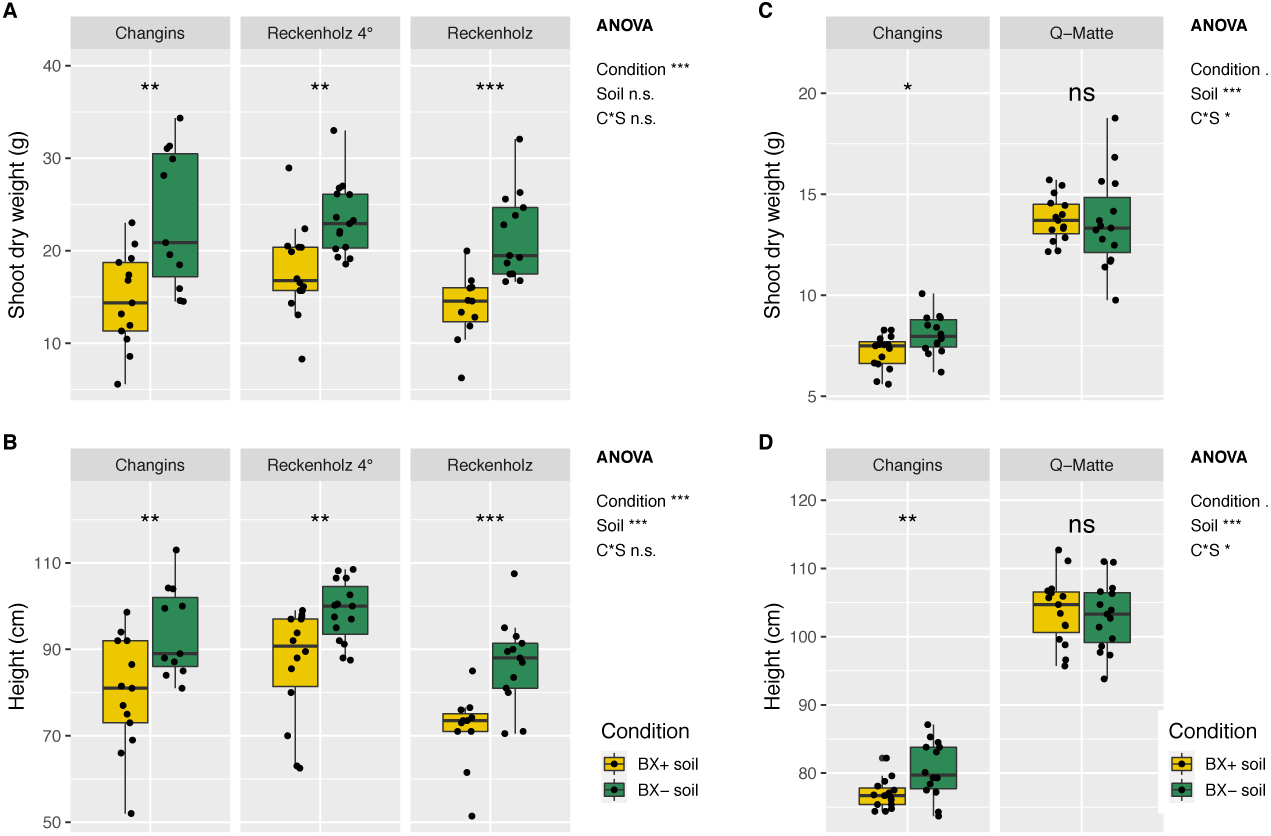
BX-feedback on maize growth. Maize plants were grown on ‘BX+’ and ‘BX-‘ variants of Changins soil as a control and compared to Reckenholz (**A**,**B**) and in a separate experiment to Q-Matte soil (**C**,**D**). The ’Reckenholz’ batch was used freshly conditioned for the feedback experiment, whereas the ’Reckenholz (4°)’ batch was store for one year in the cold room after conditioning. Shoot biomass (**A**, dry weight) and shoot height (**B**) was recorded after 10 weeks. **Data S1** documents the statistical analyses in detail. The Q-matte experiment was harvested after 9 weeks and 3 days measuring shoot biomass (**C**, dry weight) and shoot height (**D**); **Data S2** for statistic details. The ANOVA results (model: ∼ condition (C) * soil (S)) are reported next to the Figure and the pair- wise T-test results inside the panels (Significance code: *P* < 0.001 ***; *P* < 0.01 **, *P* < 0.05 *; not significant = ‘n.s.’).

%) soils when feeding on leaves of maize plants that were grown on ‘BX+’ compared to ‘BX-‘ soil variants (**Fig. 2**, **Data S1**). In the second experiment, comparing Q-Matte with Changins soil, we confirmed the feedbacks in Changins soils but we did not find feedbacks on shoot biomass (**Fig. 1C**) or height (**Fig. 1D**) for plants grown on ‘BX+’ compared to ‘BX-‘ variants of the Q-Matte soil (**Data S2**). Insect resistance could not be tested in the second experiment.

**Figure 2.**
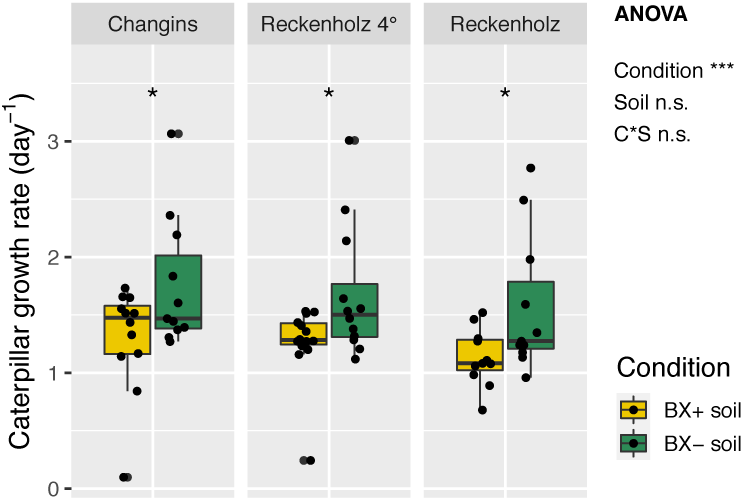
BX-feedback on maize insect resistance. Maize plants were grown on ‘BX+’ and ‘BX-‘ variants of Changins soil as a control and in two batches of Reckenholz soil. The ’Reckenholz’ batch was used freshly conditioned for the feedback experiment, whereas the ’Reckenholz (4°)’ batch was store for one year in the cold room after conditioning. Caterpillar performance of *Spodoptera frugiperda* was measured with leaves of 9 week old plants. **Data S1** documents the statistical analyses in detail. The ANOVA results (model: ∼ condition (C) * soil (S)) are reported next to the Figure and the pair-wise T-test results inside the panels (Significance code: *P* < 0.001 ***; *P* < 0.01 **, *P* < 0.05 *; not significant = ‘n.s.’).

To learn whether BX-mediated feedbacks could be preserved at low temperature, we included in the experiment the soil batch ‘Reckenholz (4°)’ which was stored for one year in the cold room. Similar to the fresh soil batch, we found significantly lower shoot biomass (**Fig. 1A**), shoot height (**Fig. 1B**) and caterpillar growth rates (**Fig. 2**) in the ‘technically overwintered’ batch (Reckenholz 4°; **Data S1**).

Taken together, BX-feedbacks on maize were in agreement with earlier findings (Hu et al., 2018), occured also in Reckenholz soil and in soil that was stored for one year in the cold room. However, no feedbacks were detected on the more fertile Q-Matte soil, revealing that BX-feedbacks are soil-dependent.

### Wheat biomass is reduced by BX-feedbacks

To answer the second major question of this study - Do BX-feedbacks also occur in wheat? - we performed different experiments to examine feedback responses (i) on growth and physiology, (ii) insect resistance and (iii) pathogen resistance. We included two wheat cultivars in most of these experiments to assess whether responses are cultivar dependent.

After having grown B73 and *bx1*(B73) maize plants in Changins and Reckenholz soils, we then tested wheat in the next generation for feedbacks. As indications for plant growth, we measured fresh and dry biomass weight of the two cultivars Drifter and Fiorina when grown on ‘BX+’ and ‘BX-‘ soil variants. Consistent with the BX-feedbacks on maize, we globally found a significantly lower fresh biomass, when wheat was grown on ‘BX+’ compared to ‘BX-‘ soil variants (**Fig. 3A**). Shoot fresh weight was reduced in both wheat cultivars and both soils (Drifter and Fiorina, - 10.1 % and -12.2 % in Changins and -9.0 % and -11.9 % in Reckenholz soil, respectively). This result was supported by factorial ANOVA but not pairwise tests (**Data S3**). This together with the observation that the BX-feedbacks were not manifested at dry biomass (**Fig. 3B**, **Data S3**) indicates that BX-feedbacks on wheat tend to be more variable and/or weaker compared to maize. Even so, we concluded that BX-feedbacks negatively impacted the growth of the two wheat cultivars in both soils similar to maize.

**Figure 3.**
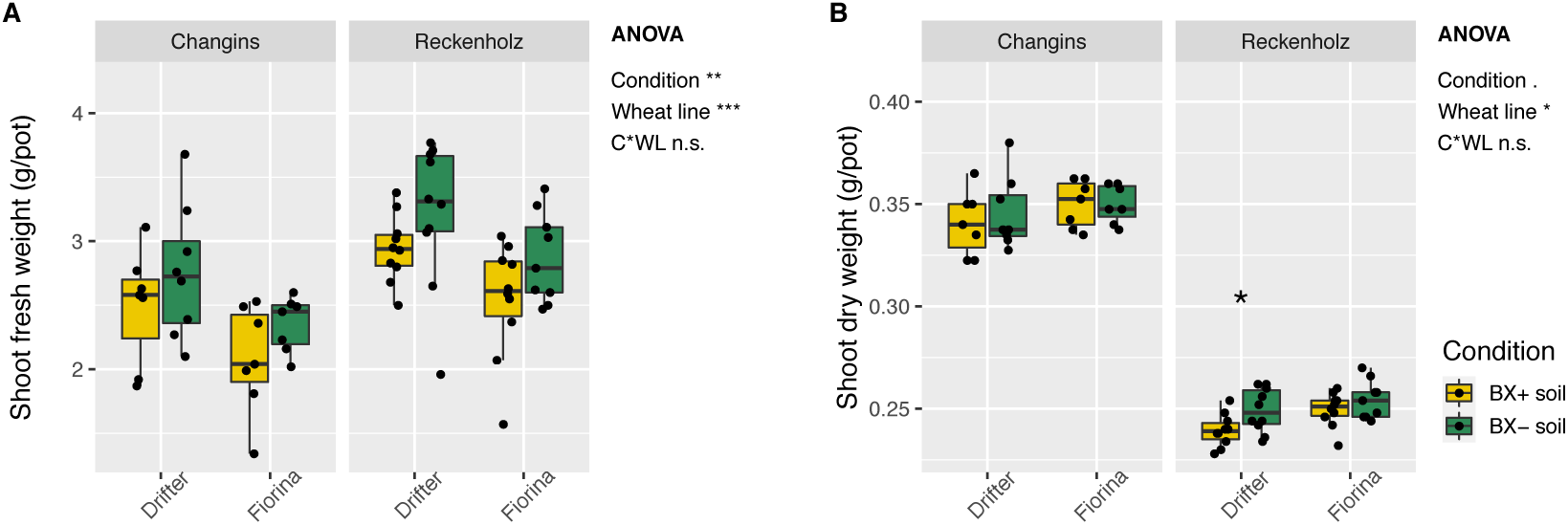
BX-feedback on wheat growth. Wheat plants were grown on ‘BX+’ and ‘BX-‘ variants of Changins and Reckenholz soils. Shoot biomass was measured as (**A**) fresh and (**B**) dry weight for the two wheat lines Drifter and Fiorina after 6 weeks of growth. **Data S3** documents the statistical analyses in detail. The ANOVA results (model: ∼ condition (C) * wheat line (WL)) are reported next to the Figure and the pair- wise T-test results inside the panels (Significance code: *P* < 0.001 ***; *P* < 0.01 **, *P* < 0.05 *; not significant = ‘n.s.’).

We also tested if BX-feedbacks affected wheat physiology and therefore, we approximated leaf chlorophyll content with SPAD measures between the third and the sixth week of wheat growth (**Fig. S1**). Although plant chlorophyll content varied more strongly with time, between the soils and between wheat varieties, we noticed a significant effect of BX-conditioning with generally higher chlorophyll content in wheat plants grown on ‘BX+’ compared to ‘BX-‘ soils (**Data S3**). This effect was most apparent in week 4 with particularly reduced chlorophyll contents in Fiorina plants grown on ‘BX-‘ soils (-20.4 % in Changins and -6.2 % in Reckenholz soil). Hence, BX-feedbacks positively affected wheat physiology by transiently enhancing leaf chlorophyll in one cultivar.

For further physiological insights, we measured the classical plant defence hormones SA, OPDA, JA, JA-Ile and ABA in plants grown on ‘BX+’ vs. ‘BX-‘ soil variants. With the exception of OPDA, hormone levels differed significantly between the cultivars with Drifter generally having higher levels of SA, JA and JA-Ile than Fiorina (**Fig. S2**, **Data S3**). For ABA, however, concentrations were higher in Fiorina compared to Drifter. Soil pre-conditioning by BXs did not affect any of the plant hormone concentrations. Thus, BX-feedbacks do not directly alter constitutive leaf hormone levels in wheat.

In summary, BX-feedbacks also occurred on wheat growth with lower fresh biomass and effects on physiology with a generally higher shoot chlorophyll content but no effects on plant defence hormone levels.

### BX-feedbacks on wheat insect resistance are cultivar-specific

In our earlier work we already found that BX-feedbacks on maize growth were genotype specific (Hu et al., 2018). Because we had studied the feedbacks of B73 plants on their B73 conditioned soils and those of W22 plants on their conditioned soils, it remained unclear whether the cause for the effects on growth occurred during the conditioning or during the feedback phase. To close this gap, we performed here an experiment where we grew B73 plants but on W22 conditioned soils. Consistent with the earlier findings in Changins soil, we measured also in potting soil a significantly lower shoot biomass (**Fig. 4A**) and lower insect performance (**Fig. 4B**) for the B73 response plants on their own conditioned ‘BX+’ and ‘BX-‘ soil variants (**Data S4**). When growing B73 as response plants on the soil variants conditioned by W22 lines, we found the same negative growth and insect phenotypes as on B73 conditioned soils (**Fig. 4**) while this was not the case for W22 as response genotype (Hu et al., 2018). Hence, we identified the feedback and not the conditioning phase as causal and we concluded that the different genotypes of the response plant explain the different findings.

**Figure 4.**
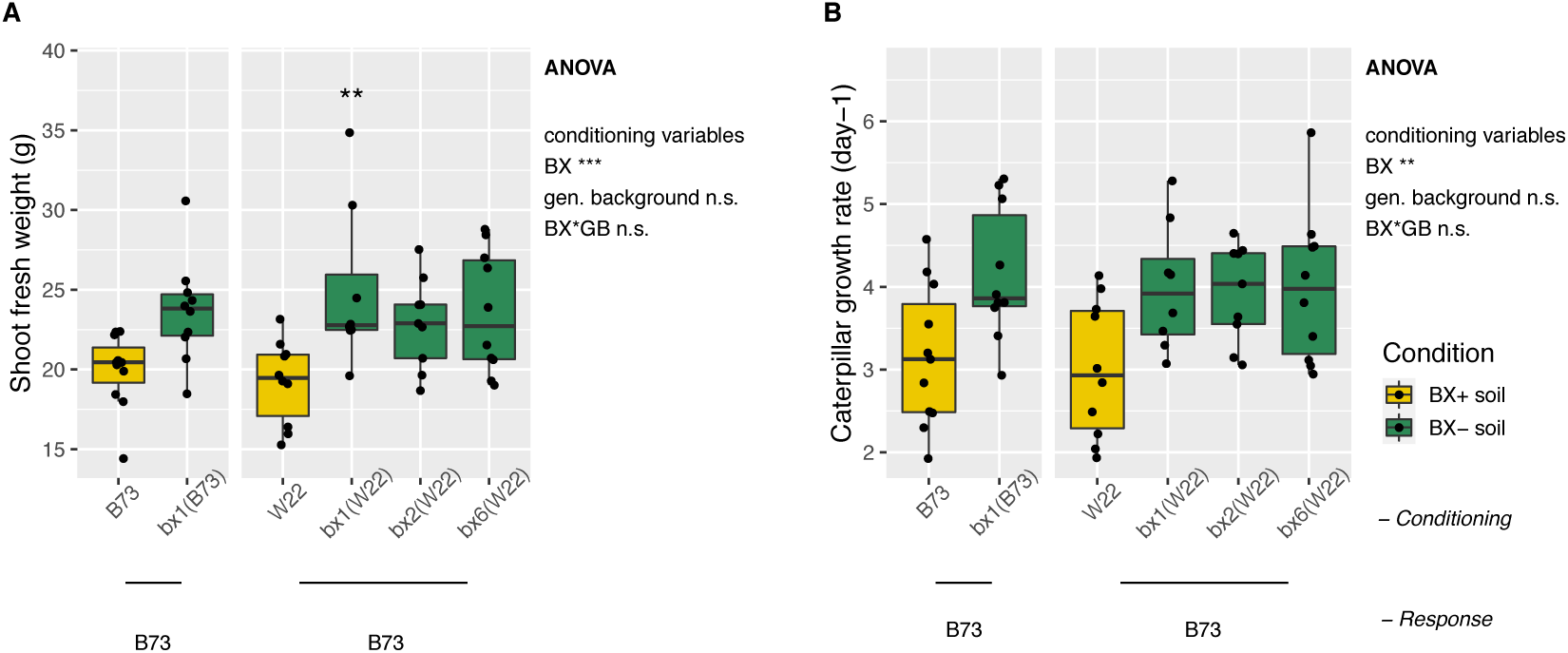
Genetics of BX-feedback on maize. Potting soil was conditioned with B73 and *bx1*(B73) as well as with W22, *bx1*(W22), *bx2*(W22) and *bx6*(W22) followed by a feedback phase with only B73 plants (n = 8–11). Ten week old plants were utilized for measuring (**A**) shoot biomass (fresh weight) and (**B**) *Spodoptera frugiperda* performance. **Data S6** documents the statistical analyses in detail. The ANOVA results (model: ∼ BX condition (BX) * genetic background condition (GB)) are reported next to the Figure and the pair-wise Tukey-test results inside the panels (pair-wise comparison with wild-type, significance code: *P* < 0.01 **, *P* < 0.05 *; not significant = ‘n.s.’).

As the genotype dependent BX-feedbacks on maize specifically enhanced insect resistance, we also examined the two wheat cultivars Drifter and Fiorina for eventual BX-feedbacks when grown on ‘BX+’ and ‘BX-‘ soil variants. Plant insect resistance was assessed indirectly by measuring growth rates of the generalist caterpillar *S. littoralis* feeding wheat leaves. ANOVA analysis indicated that the two cultivars behaved differently in the two soil variants (**Data S5**). While insect growth rates were unaffected by BX-conditioning in Drifter, they were significantly higher on Fiorina when growing on ‘BX+ soil’ (**Fig. 5A**). This suggested that BX-feedbacks reduced the insect resistance of the wheat cultivar Fiorina while Drifter, the cultivar with generally higher defence hormone levels, was unaffected in its insect resistance.

**Figure 5.**
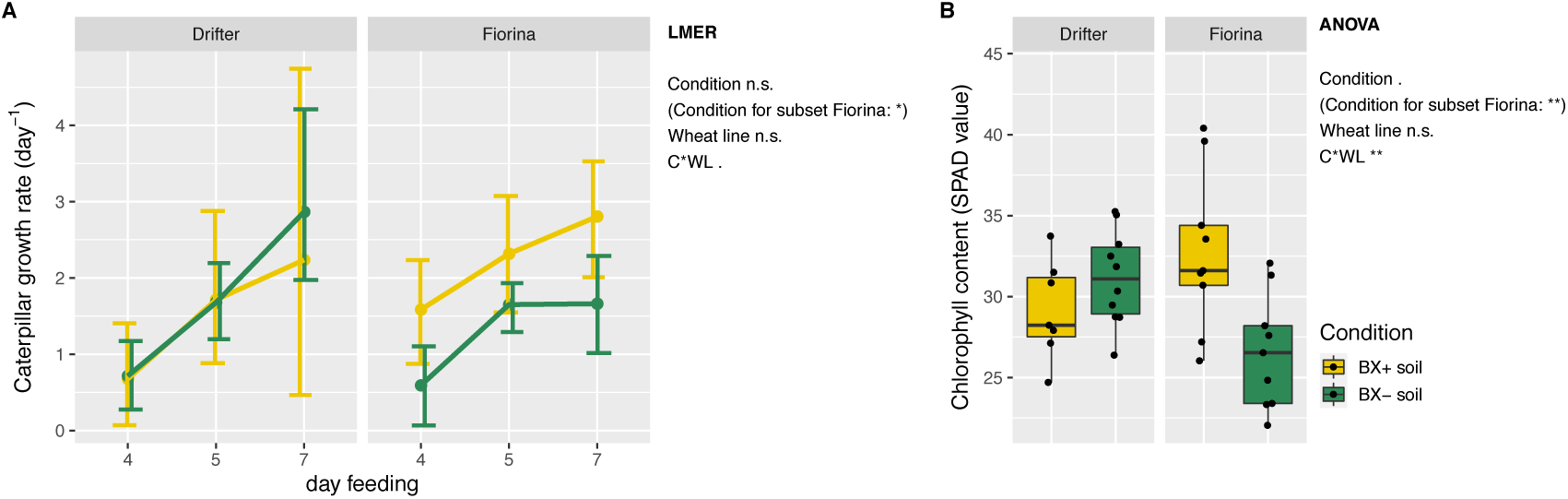
BX-feedback on wheat insect resistance and chlorophyll content. Wheat plants from cultivar Drifter and Fiorina were grown on ‘BX+’ and ‘BX-‘ variants of Changins soil. (**A**) Caterpillar performance of *Spodoptera littoralis* fed with leaves of 5 weeks old plants was measured after 4, 5 and 7 days of feeding; and (**B**) leaf chlorophyll content (SPAD values) was measured on 6 weeks old plants. **Data S4** documents the statistical analyses in detail. The LME (model: ∼ day_feeding * condition (C) * wheat_line (WL)) for (**A**) and ANOVA (model: ∼ condition (C) * wheat_line (WL)) for (**B**) results are reported next to the figure (significance code: *P* < 0.001 ***; *P* < 0.01 **, *P* < 0.05 *; ‘n.s.’ = not significant).

To confirm the BX-feedbacks on leaf chlorophyll content in the previous experiment, we repeated the SPAD measurements. Consistently with the previous results (**Fig. S1**) we found again significantly reduced chlorophyll contents (-20.8%) in Fiorina plants when grown on ‘BX-‘ soils (**Fig. 5B**, **Data S5**). And, as indicated by the significant interaction term of the ANOVA, there was no effect on Drifter plants. It appears that enhanced chlorophyll content, as a positive BX-feedback, presents a cultivar-specific response.

Overall, we conclude that BX-feedbacks on wheat are cultivar-specific as evidenced by the insect resistance and shoot chlorophyll assays.

### No detectable impact of BX-feedbacks wheat pathogen resistance

To evaluate whether BX-conditioning would also affect wheat pathogen resistance, we tested feedbacks to inoculations with *Zymoseptoria tritici*. We scored pycnidia, which are the reproductive structures attesting successful fungal multiplication in the host tissue. There was no significant difference in pycnidia counts of plants growing on ‘BX+’ or ‘BX-‘ soils (**Fig. S3A**, **Data S6**). Plants growing on ‘BX+ soil’ tended to have higher numbers of pycnidia compared to plants growing on ‘BX-‘ or the control soil). We also quantified *Z. tritici* in the infected leaf using qPCR, but pathogen abundance did not differ neither between the two BX conditions nor to the control soil (**Figure S3B, Data S6**). Taken together, we found no evidence that the BX-feedbacks impact the resistance of the wheat cultivar Drifter to the pathogen *Z. tritici*.

## Discussion

BX-mediated microbial feedbacks influenced maize growth and insect resistance in one soil (Hu et al., 2018). Here, we explored these BX-feedbacks more broadly, asking first, whether they also occur in different soils and secondly, whether they also influence wheat as a typical crop following maize in European rotation schemes. We performed the latter experiments with two wheat cultivars, because our earlier maize work had revealed genetic differences in responsiveness to BX-feedbacks.

### BX-feedbacks are soil dependent

Consistent with our earlier work in soil from Changins (Hu et al., 2018), we also find BX-feedbacks on maize growth and insect resistance in the tested Reckenholz soil (**Figs. 1 & 2, Data S1**). The feedbacks were of similar strength and in the same direction with reduced shoot biomass, lower shoot height and enhanced insect resistance for maize plants grown on ‘BX+’ compared to ‘BX-‘ soil variants. Using sterilization we had demonstrated earlier that the differential feedbacks of maize growing either on ‘BX+’ and ‘BX-’ variants of the Changins soil were driven by the microbiota (Hu et al., 2018). We had also shown that the root and rhizosphere bacterial and fungal communities of the maize plants, which conditioned these soil variants (WT -> ’BX+’, *bx1* -> ‘BX-‘), differed in their composition. Therefore, it is tempting to assume that BX exudation also changes the microbiota in the Reckenholz soil and that different microbiotas in ‘BX+’ and ‘BX-’ soil variants explain the differential feedbacks. While we have no experimental proof for the second part of the assumption, we found differences in community composition analysing the root and rhizosphere microbiotas of the maize lines, which conditioned the ’BX+’ and ‘BX-‘ variants of the Reckenholz soil, too (Cadot et al., 2020). Of note, these microbiota analyses were performed on the exact same plants, which conditioned the ’BX+’ and ‘BX-‘ variants of the Reckenholz soil tested in this study. In Cadot et al. (2020) we investigated the general structuring of BX exudation on the root and rhizosphere microbiota and we found little overlap among BX-sensitive microbes in different field soils. This implies that different sets of BX-sensitive microbiotas can trigger similar feedback effects and that there is a functional overlap among them in different soils.

With this study we now know that BX-feedbacks on maize function in the clay loam soil Changins and the loam soil Reckenholz, but not the silt loam Q- Matte soil. As the maize plants grew best in this soil (**Fig. 2**), we consider Q-Matte as the most fertile among the tested soils. Given the evidence that BX-feedbacks lose their effectivity at higher soil fertility, further work specifically investigating the impact of soil fertility on BX-feedbacks is required. To what extent the feedbacks are affected by different soil histories (arable vs. grassland) also remains to be determined. Finally, further work is necessary to clarify the range of effective BX-feedbacks across various soil types and soil texture classes. A systematic examination with conditioning ‘BX+’ and ‘BX-‘ variants in fields of different soil types will present a tremendous effort, diverse soil types could be collected and then conditioned in the greenhouse to work out the impact of specific biogeochemical properties on BX-feedbacks. Ultimately, understanding the context dependency of such BX-feedbacks appears important, hypothesising that these feedbacks do not only occur under controlled greenhouse but also under realistic agricultural conditions.

### BX-feedbacks are preserved at 4°C for at least one year

We knew that BX-feedbacks are preserved over a winter period in the field (Hu et al., 2018). We had left overwintering the ‘BX+’ and ‘BX−‘ soil variants after conditioning them in summer and had observed the feedbacks still in the following spring. Prompted by this observation and motivated to evaluate our experimental approach, we wanted to answer, if ‘BX+’ and ‘BX−‘ soil variants can be stored in a cold room at 4°C for at least one year without losing their efficacy. Similar to the field winter, we found that a ‘technical’ overwintering of the ‘BX+’ and ‘BX−‘ soil variants in the cold room also preserved the BX-feedbacks (**Figs. 1** & **2, Data S1**). This retention of feedback capacity has important practical consequences as there is no need to study BX-feedbacks immediately on freshly conditioned soils and that batches of conditioned soil can be stored and examined at later stages. It will be of obvious interest to study further for how long BX-feedbacks can be preserved both technically in the cold room but also in the field and whether the strength of the effects decreases over time?

There is evidence for persistence of PSFs longer than one plant generation. For instance, *Jacobea vulgaris*, a plant species of the Asteraceae family, was still responsive in the third generation to a first conditioning phase despite the growth of a second conditioning phase by different plant species in these soils (Wubs and Bezemer 2018). Longer term effects were seen with microbiota legacies inherited through different plant community diversities growing for 11 (Schmid et al., 2019) or 8 years (Schmid et al., 2021). Because PSFs can persist over multiple generations, it is possible that BX feedbacks are effective later points in time, i.e. later than an overwintering period (Hu et al., 2018) or later than one year as shown with storage in the cold room (**Fig. 1**). Understanding the long-term persistence of BX-feedbacks appears also relevant with regard to agriculture where eventual feedbacks may persist in the crop rotation.

### BX-feedbacks also function on wheat

Like maize, wheat also produces BXs (Li et al., 2018; Niemeyer, 2009) and secretes them to the rhizosphere (Chen et al., 2010). Although a broad examination how other or non-BX producing plant species such as for instance functional groups like legumes, forbs or crucifers deal with BX-conditioned soils would have been interesting, we chose to investigate wheat, as it is often following maize in crop rotations in Europe and because of agronomic relevance.

From our study, it is clear that BX-feedbacks also function on wheat. Similar to maize, negative feedbacks on biomass were seen when wheat was grown on ‘BX+’ compared to ‘BX-‘ soil variants (**Fig. 3, Data S2**). Although not compared side-by-side, it appeared that BX-feedbacks on wheat growth tended to be more variable and/or weaker compared to maize. BX-feedbacks were also observed on wheat physiology (**Figs. S1 & 4B**, **Data S1** and **S3**) and insect resistance (**Fig. 4A**), but the direction of the feedbacks contrasted with maize responses to soil BX-conditioning. While the chlorophyll approximations were higher on ‘BX+’ soil in the tested wheat cultivars, they were lower in maize (Hu et al., 2018). Opposite effects were also seen on insect performance with enhanced resistance of maize grown on ‘BX+’ soil (Hu et al., 2018), while the wheat cultivar Fiorina was more susceptible when grown on ‘BX+’ soil. At present such opposite BX feedback behaviours cannot be generalized for being either maize or wheat specific, not only because more cultivars per plant species would be required but mainly because both plant species exhibited strong genetic variation in their feedback responses (see below).

### Microbial and allelopathic components of BX-feedbacks

‘BX+ soils’ contain both, BX chemicals and a thereby conditioned microbiota (‘BX+ microbiota’), while the BX compounds are absent in ‘BX- soils’ and therefore, the microbiota is not conditioned by BXs (‘BX- microbiota’). Thus, the differential feedbacks observed on plants have a microbial (BX+ vs. BX- microbiota) and a chemical component (presence vs. absence of BXs). With our maize work, we had demonstrated the presence of this microbiota component using sterilization and complementation experiments (Hu et al., 2018). Sterilization did not affect the MBOA (being the most abundant BX) levels in BX+ soils, but abolished the feedbacks, which were restored with the addition of a microbial extracts to the sterilized and MBOA containing soils. We had formulated a model based on the presence of BXs and the microbiota presenting the drivers for the BX feedbacks. For the maize line B73, with which the sterilization and complementation experiments were performed, it is clear that feedbacks have a microbial and not an allelopathic component. Nevertheless, we explicitly include in the model the possibility of direct contributions of BX chemicals to the feedbacks, in particular, because BXs are renowned for their allelopathic effects on many plant species (Schandry and Becker, 2020).

Because of the following two evidences, we speculate that the relative contributions of microbial and allelopathic components in BX feedbacks differ among response species or genotypes. For instance, the reductions in plant biomass of both maize (**Fig. 1**) and wheat (**Fig. 3**) intuitively suggest an allelopathic component in this feedback, as such effects of BX on cereals has been demonstrated (Acharya et al., 2020). Nevertheless, for maize that grows on the maize conditioned BX+ soils, the sterilization and complementation demonstrated clear a microbial and not an allelopathic one in the feedbacks (Hu et al., 2018). The previous conditioning of the soil by maize resulted in an enrichment of maize-adapted microbes in the BX+ soils and an open question is how wheat responds to these maize-adapted microbes. Therefore, it remains to be shown for wheat whether the negative growth feedback is also consequence of microbial interactions or if this is not at least partly an allelopathic effect. Complementary experiments utilizing a BX-defective wheat mutant (currently not available) that produces BX+ soils containing wheat-adapted microbes and performing reciprocal feedback experiments with maize would be necessary for understanding the role of microbial adaption in BX-feedbacks.

The second evidence, why we think of microbial and allelopathic components in BX feedbacks, is related to microbiota perception by the response plants. Maize plants growing on BX+ soils exhibited elevated defence hormone levels and expression of defence marker genes that are reminiscent of ISR (Hu et al., 2018) and therefore, this point to a microbial component in the BX feedbacks. In contrast, wheat did not respond with altered defence hormone levels suggesting no microbial contribution to the feedbacks. This interpretation could be further corroborated with experiments testing for eventual microbiota differences of wheat when growing on maize conditioned BX+ or BX- soils. As indicated above, the associations and thereby the feedbacks to a maize conditioned microbiota in BX+ soils may differ between maize or wheat as host plants in the subsequent generation. A hypothesis is that the host plant in the next generation and the previously conditioned microbiota (here by maize) need to be adapted to each other to express a microbiota induced systemic resistance.

Taken together, dedicated work related to microbiota adaptation and plant responsiveness is now needed to disentangle the microbial from allelopathic contributions to the observed BX-feedbacks.

### Genetic variation in responsiveness to BX-feedbacks

Insect resistance and chlorophyll content were unaffected in the cultivar Drifter, whereas Fiorina was more susceptible to the *Spodoptera littoralis* and had reduced chlorophyll levels when grown on ‘BX+ soil’ (**Fig. 4**). Similar to the genotypic variation we found in the tested two wheat cultivars to BX-conditioned soils in this study, the two maize lines B73 and W22 expressed both the BX- dependent increased insect defences whereas growth suppression was only seen in B73 but not in W22 (Hu et al., 2018). Here, we now demonstrated that the genotype of the response plant explains the different growth feedbacks (**Fig. 4**). It is not uncommon that different plant genotypes express differential responsiveness to microbes. Wheat cultivars respond differently to associations with beneficial microbes (Akbari et al., 2020; Egamberdieva, 2010; León et al., 2020) as well as express different sensitivities to the allelopathic effect of BXs (Schulz et al., 2013). In the context of microbially triggered ISR, there is genetic variation in responsiveness as reported for barley (Shrestha et al., 2019). This study showed that an appropriate genetic background is required to perceive the microbial priming in order to express enhanced resistance. The common conclusion is obvious: there is genetic variation in host plant responsiveness to individual microbes or complex soil microbiomes as seen also in the BX feedbacks. The underlying mechanisms, how plants perceive a specific complex soil microbiota that induces systemic resistance or promoting growth, are not know. Ultimately, plant loci for positive responsiveness to microbiota feedbacks will open new opportunities to integrate beneficial plant-microbiome interactions into crop breeding programs.

### Relevance of BX-feedbacks in agriculture

Similar to Pineda et al. (2019), our studies on BX-feedbacks serve as evidence that the concept of PSFs may be exploited in cropping systems for pest control. There is often a discrepancy between results obtained in highly controlled greenhouse experiments and those in real-life field trials (Beals et al., 2020; Brinkman et al., 2010; Forero et al., 2019). Nevertheless, several findings of our work under controlled conditions argue that these BX feedbacks may be relevant for agriculture: (i) BX-feedbacks were observed in natural field soils, (ii) BX-feedbacks preserve their effectivity over a winter period and (iii) agronomically relevant crops respond to BX-feedbacks. Field experiments with natural conditions and a rotation of wheat following maize are now necessary to answer the major emerging question from this work: do these BX-feedbacks have real-life agronomic implications?

## Authors’ contributions

KS, ME and MvdH conceived the original project and KS supervised the experiments. ASV, DC, ME, and MvdH provided field resources, laboratory infrastructure and seeds. SC, VG and LH performed the experiments with support by GG for analytical work, by NS for Zymoseptoria work and support by ASV for qPCR analysis. SC and KS analysed the data. SC and KS wrote the manuscript with input from all authors.

## Acknowledgements

We thank Damian Amrein from the seed selection group (Agroscope) for providing the Fiorina wheat seeds. Special credits go to Nicolas Widmer, Pierre Pignon, Juerg Hiltbrunner and Fritz Kaeser (Agroscope) for field support and Philip Streckeisen (Agroscope) for technical support in the greenhouse. We also thank Diane Buerge (Agroscope) and Dr. Moritz Bigalke (Institute of Geography, University of Bern) for soil analyses. qPCR experiments were performed at the Genetic Diversity Centre (GDC), ETH Zurich.

## Supplementary Information

### Supplementary Methods

#### Soil conditioning

##### Field experiment ‘Reckenholz 2015’

Several batches of conditioned soils were either prepared in field or in greenhouse experiments (**Table S1**). The conditioned soil batch ‘Reckenholz 2015’ was harvested from a field experiment, which we performed in 2015 with the same setup as the field experiment in 2016. We grew single B73 and *bx1*(B73) plants separated by maize hybrids in a field at Agroscope. We used parcel 212 in 2015, whereas in 2016 it was parcel 209 (80 meters apart). Field preparation included, harrowing (April 24th) and seeding of the hybrid plants (May 19th). Similar to the experiment ‘Reckenholz 2016’, on June 5^th^, we replaced individual hybrid plants and transplanted pre-grown B73 and *bx1*(B73) to the field. The setup were two transects (orthogonal to the rows of maize hybrids, spaced by 1 m with 5 to 7 hybrid plants between test alternating plants). Field management followed conventional farming practices consisting of herbicide applications (June 11^th^: 1.5 l/ha of Laudis (44 g/l Tembotrione and 22 g/l Isoxadifen-ethyl) and 2 l/ha Aspect (333 g/l Terbuthylazin, 200 g/l Flufenacet) and mineral fertilizer applications of 2.5 dt/ha of ammonium nitrate (27%) and magnesium (2.5%), 1.90 dt/ha (June 4th) and 1 dt/ha of urea (June 24^th^). Preceding crop was artificial pasture (2013-2014). Following the common protocol (Cadot et al., 2020; Hu et al., 2018): after three months of growth, the shoots of the maize plants were cut, a 20 x 20 x 20 cm soil core was dug out around the maize plants, the root system removed, and the soil stored at 4°C until use. The particularity of the collected soil cores from the ‘Reckenholz 2015’ batch is that they were stored then for one year at 4°C in the cold room before use.

##### Soil conditioning in the greenhouse (for Experiment 2)

For Experiment 2, field soils were collected at Agroscope in Changins (parcel 29, 46° 23’ 58.468’’, N 6° 14’ 23.008’’ E; between the rows of winter wheat seedlings) and on a grassland meadow named ‘Q-Matte’ close to Bern (46° 57’ 20.465799’’ N, 7° 19’ 59.067782’’ E). The soils were sieved to 10 mm, homogenized and filled into 2L pots (Rosentopf Soparco, Hortima). For soil conditioning the plants were grown under controlled conditions (day/night: 16 h/8 h; temperature: 26 °C/23 °C; light: 550 µmol m^-2^s^-1^; humidity: 50%) in a walk-in climate chamber for 6 weeks. Two and four weeks after sowing plants were fertilized with 100 ml of a nutrient solution (0.2% [w/v]; Plantactive Typ K, Hauert) supplemented with iron (1 ‰ [w/v]; Sequestrene rapid, Maag). Plants were watered as needed and weekly randomized.

##### Soil conditioning in the greenhouse (for Experiments 3 to 5)

Additional Changins soil was conditioned in the greenhouse for the wheat experiments 3 to 5 (**Table S1**). For this, a large bulk of unconditioned field soil was collected at Agroscope in Changins (parcel 30-31, 46° 24’ 0.792’’ N 6° 14’ 23.531’’ E). To condition the soil, maize was grown between the 28.05.2017 and the 24.08.2017 in 3L pots with controlled conditions (8:00 to 22:00 light period, 26°C ± 2 °C, 55% relative humidity). The plants were fertilized every week with NPK (99/99/74 g/l) and micronutrients (Wuxal Universal, Hauert; 20 mL/10L water) and with iron supplement (Sequestrene rapid, Maag; 5.5% Fe in the form of EDDHA, 20g/10L water). Plants were uniformly watered with tap water when needed. Replicate samples of conditioned soils were similarly collected as in the field experiments: conditioning was stopped after 3 months, the maize shoot and the root system removed, the whole soil of a 3L-pot was harvested as a replicate of conditioned soil and stored at 4°C until use.

##### Detailed setups of the individual feedback experiments

###### Experiment 1 – BX-feedbacks on maize in Reckenholz soil

The first feedback experiment was performed with maize in the greenhouse following a fully randomized design testing the factor *conditioning* (BX+ or BX-) in 3 soil batches. Three field-conditioned soil batches included ‘Changins’ as a control soil and two batches of Reckenholz soil (**Table S1**). The batch ‘Reckenholz’ was freshly conditioned in 2016 while ‘Reckenholz (4°)’ was collected in 2015 and then stored for one year at 4° until use. For each combination of soil batch and BX condition, ten 3L-pots were prepared and each seeded with one B73 maize plant resulting in a total of 60 experimental units (3 soil batches * 2 BX conditions * 10 replicates; see **Data S1** for experimental design). The plants were grown in the greenhouse using the following conditions: 26°C ± 2 °C, 55% relative humidity, 8:00 to 22:00 light period. Fertilization was also brought every week with the watering, with NPK (99/99/74 g/l) and micronutrients (Wuxal Universal, Hauert; 20 mL/10L water) and with iron supplement (Sequestrene rapid, Maag; 5.5% Fe in the form of EDDHA, 20g/10L water). Plants were uniformly watered with tap water when needed. At 9 weeks the first fully expanded leaf was utilized for caterpillar assay. At 10 weeks, plant height was measured and the remaining aboveground biomass harvested and weighed after drying in an oven at 60°C for 72h.

###### Experiment 2 - BX-feedbacks on maize in Q-Matte soil

To further test BX-feedbacks we conducted a second experiment with greenhouse-conditioned batches of ’Q-Matte’ soil along with ‘Changins’ soil as a control (**Table S1**). The experimental design was fully randomized and full factorial (2 soils * 2 BX condition; BX+ or BX-) with 15 replicates (total of 60 experimental units; see **Data S2**). Two litre pots containing the differentially conditioned soils were seeded with B73 maize plants and grown under controlled conditions (day/night: 16 h/8 h; temperature: 26 °C/23 °C; light: 550 µmol m^-2^s^-1^; humidity: 50%) in a walk-in climate chamber. Plants received 100 ml fertilizer (0.2% [w/v] Plantactive Typ K (Hauert) supplemented with 1 ‰ [w/v] Sequestrene rapid (Maag)) at two, four and six weeks and were watered as needed and weekly randomized. The experiment was stopped at 9 weeks and 3 days of growth, measuring plant height and harvesting maize shoots, which were dried (80 °C for 72 h) and weighed on a microbalance.

###### Experiment 3 – BX-feedbacks on wheat on growth and physiology

The third feedback experiment was performed with wheat, again, with a randomized design testing the factor *conditioning* (BX+ or BX-). The experiment was conducted with a greenhouse-conditioned batch of Changins soil and a field- conditioned batch of Reckenholz soil (**Table S1**). Because of fewer amounts of the Changins soil, this part of the experiment was performed in 0.5 L pots (16 experimental units = 2 BX conditions * 8 replicates; see **Data S3** for experimental design). The other part of the experiment with Reckenholz soil was conducted with 1L pots (20 experimental units = 2 BX conditions * 10 replicates; **Data S3**). Because of the differences in pot size and how the soils were conditioned, we do not compare the two soils in this experiment. We grew the two wheat cultivars ‘Drifter’, a hard winter wheat cultivar, and ‘Fiorina’, a spring wheat currently used by Swiss farmers. Six seeds were sowed per pot and two weeks after seeds germinated the number of plants was reduced to 5 for plants growing in the bigger pots, and reduced to 4 for the ones in smaller pots. Growth conditions in the greenhouse included a light period from 8:00 to 22:00, 70% relative humidity and temperatures of 22°C (day) and 18°C (night). Fertilization was done by mixing NPK granules (Osmocote exact, Standard 5-6M; 15-9-12 NPK, 2MgO and micronutrient; 4g / kg soil) into the soils before filling the pots. Plants were uniformly watered with tap water when needed. Chlorophyll content was measured at 3, 4, 5 and 6 weeks after sowing. The experiment was harvested at 6 weeks (plants were before flowering) as follows: the second fully unfolded leaves were cut, immediately wrapped in aluminium paper, shock-frozen in liquid nitrogen and then stored at -80° until hormone analysis. The remaining aboveground shoots were cut and recorded as fresh and dry weight (after drying in an oven at 60°C for 72h ) biomass per pot.

###### Experiment 4 – BX-feedback on wheat insect resistance

A fourth experiment, using a fully randomized design for testing the factor *conditioning* (BX+ or BX-), was set up in a climate chamber to examine insect performance. Experiment 4 was conducted with a greenhouse-conditioned batch of Changins soil (**Table S1**). The conditioned soils (BX+ and BX- variants) were further sieved to 5 mm, filled in 300 mL pots and sown with four wheat plants. We used again the two wheat cultivars ‘Drifter’ and ‘Fiorina’ resulting in 40 experimental units (2 BX conditions * 2 cultivars * 10 replicates; **Data S4**). Plants were grown with a light period from 8:00 to 22:00 at 22 °C and 18°C at night, with 70% relative humidity. At five weeks, when we terminated the experiment, Drifter was at the vegetative stage while Fiorina started flowering. We approximated the chlorophyll content and used two fully expanded leaves per plant for a caterpillar performance assay.

###### Experiment 5 – BX-feedbacks on wheat pathogen resistance

We examined wheat resistance against *Zymoseptoria tritici* in a fifth experiment, again with a fully randomized design testing the factor *conditioning* (BX+ or BX-). A greenhouse-conditioned batch of Changins soil was employed for this climate chamber experiment (**Table S1**). With the goal to prepare a control soil we sterilized BX+ and BX- soil variants with x-rays. The BX+ and BX- soils were prepared by re-inoculating to them with microbial washes (Hartman et al., 2017) extracted from non-sterilized BX+ and BX- soils. A 1:1 mixture of sterile BX+ and BX- soils without microbial wash was finally used as control soil. The experiment was set in three blocks (so that plants could be sampled at three timepoints) and ten replicates resulting in 90 experimental units (3 soil variants * 3 timepoints * 10 replicates; **Data S5**). These soils were filled into 200 mL pots and planted with four Drifter seeds per pot, and then cultivated at 18°C, 70% relative humidity for 3 weeks until starting the pathogen assay.

###### Experiment 6 – Genetics of BX-feedbacks on maize

The Experiment 6 was performed with maize using regular potting soil (Klasmann–Deilmann GmbH, Germany), which was conditioned in 3L-pots (13.1 cm depth and 20.2 cm diameter) in the greenhouse. The conditioning was not only done with B73 and *bx1*(B73) but also with the maize lines W22, *bx1*(W22), *bx2*(W22) and *bx6*(W22). The growth conditions and maize lines were described earlier (Hu et al., 2018; ’Benzoxazinoid pathway experiment’). Different compared to the earlier experiment is that here we utilized potting soil and we only grew B73 plants on B73 and W22 conditioned soils (n = 8–11 replicates; see **Data S6** for experimental design). Pots were randomly placed on a greenhouse table (26 °C ± 2 °C, 55% relative humidity, 14:10 h light/dark, 50,000 lm m^−2^) and re-arranged weekly. Plants were watered three times per week. Ten weeks after planting, the shoot biomass and larval growth were measured.

## Supplementary Figures

**Figure S1.**
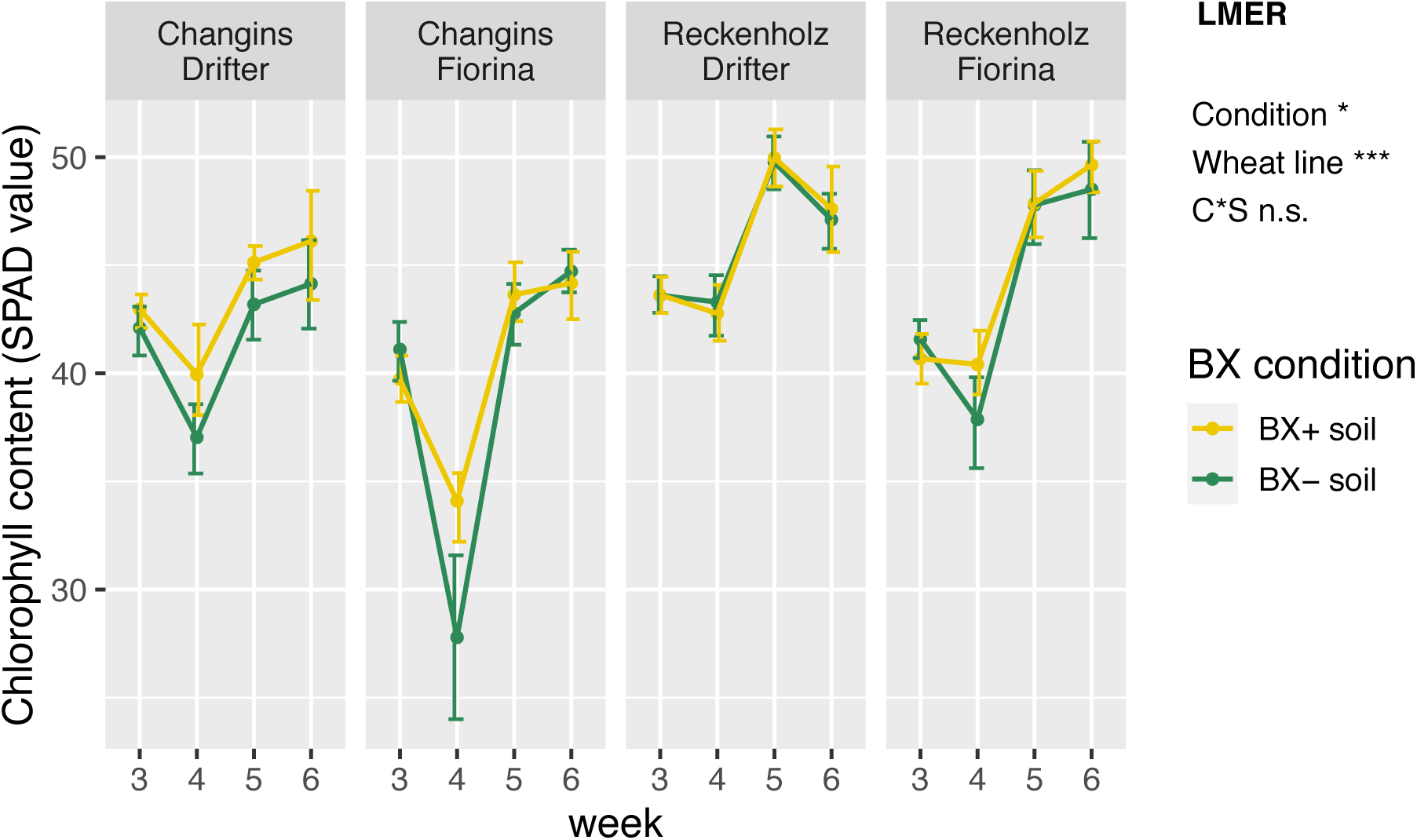
BX-effect on wheat leaf chlorophyll content during growth. Wheat plants from cultivar Drifter and Fiorina were grown on ‘BX+’ and ‘BX-‘ variants of Changins and Reckenholz soil. Mean chlorophyll content (SPAD values) per pot was measured from the 3rd to the 6th week of plant growth. Data S3 documents the statistical analyses in detail. The LME results for condition and wheat line variables (model: ∼ week * soil and pot size * condition (C) * wheat line (WL)) are reported next to the figure (significance code: *P* < 0.001 ***; *P* < 0.01 **, *P* < 0.05 *; ‘n.s.’ = not significant).

**Figure S2.**
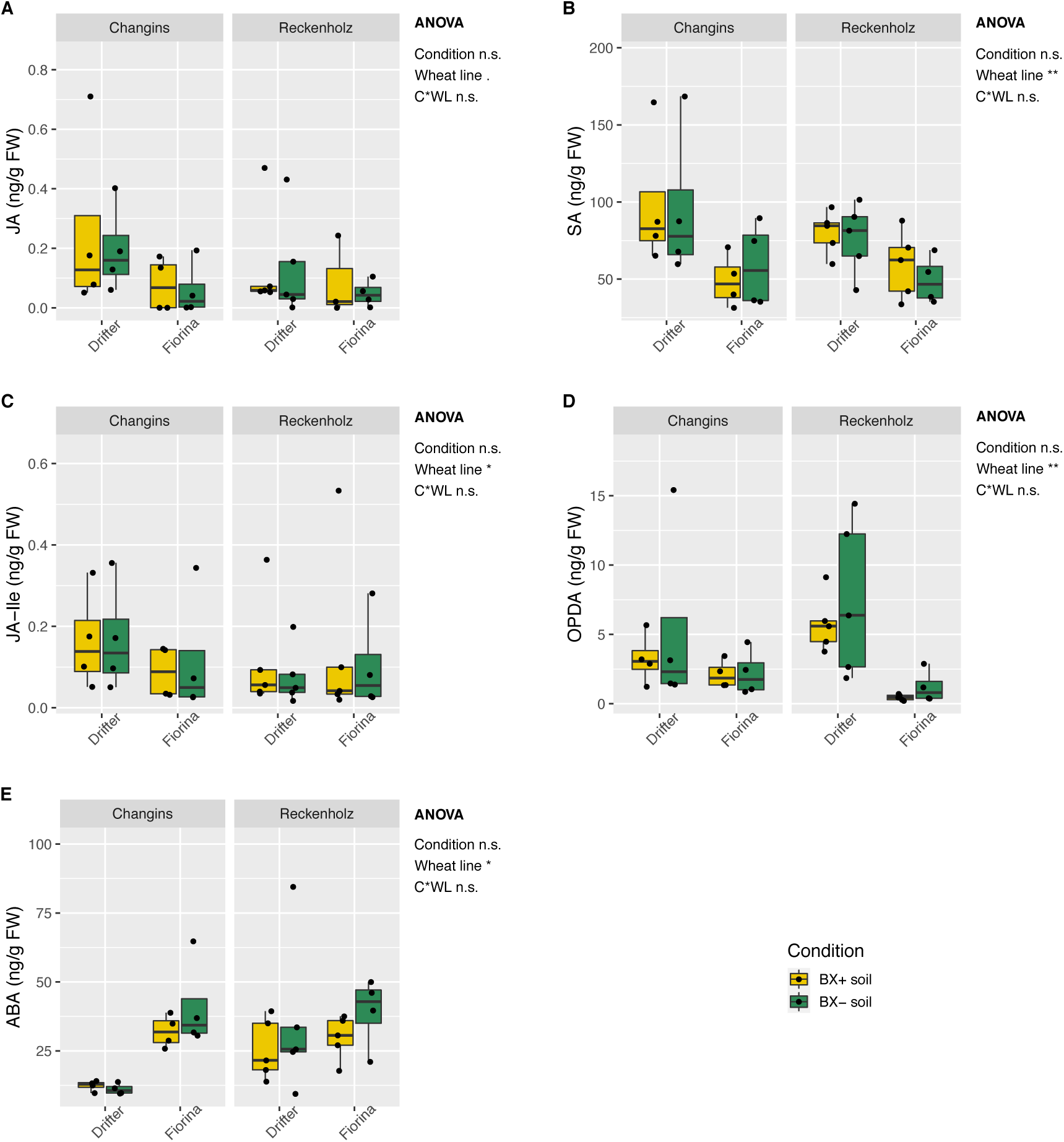
BX-effect on wheat hormone concentration. Wheat plants from cultivar Drifter and Fiorina were grown on ‘BX+’ and ‘BX-‘ variants of Changins and Reckenholz soil. Leaf concentration of (A) jasmonic acid (JA), (B) salicylic acid (SA), (C) jasmonic acid – isoleucine (JA-Ile), (D) oxophytodienoic acid and (E) abscicic acid (ABA) were measured in 6 week old plants. Data S3 documents the statistical analyses in detail. The ANOVA results for condition and wheat line (model: ∼ soil and pot size * condition (C) * wheat line (WL)) are reported next to the figure (significance code: *P* < 0.001 ***; *P* < 0.01 **, *P* < 0.05 *; ‘n.s.’ = not significant).

**Figure S3.**
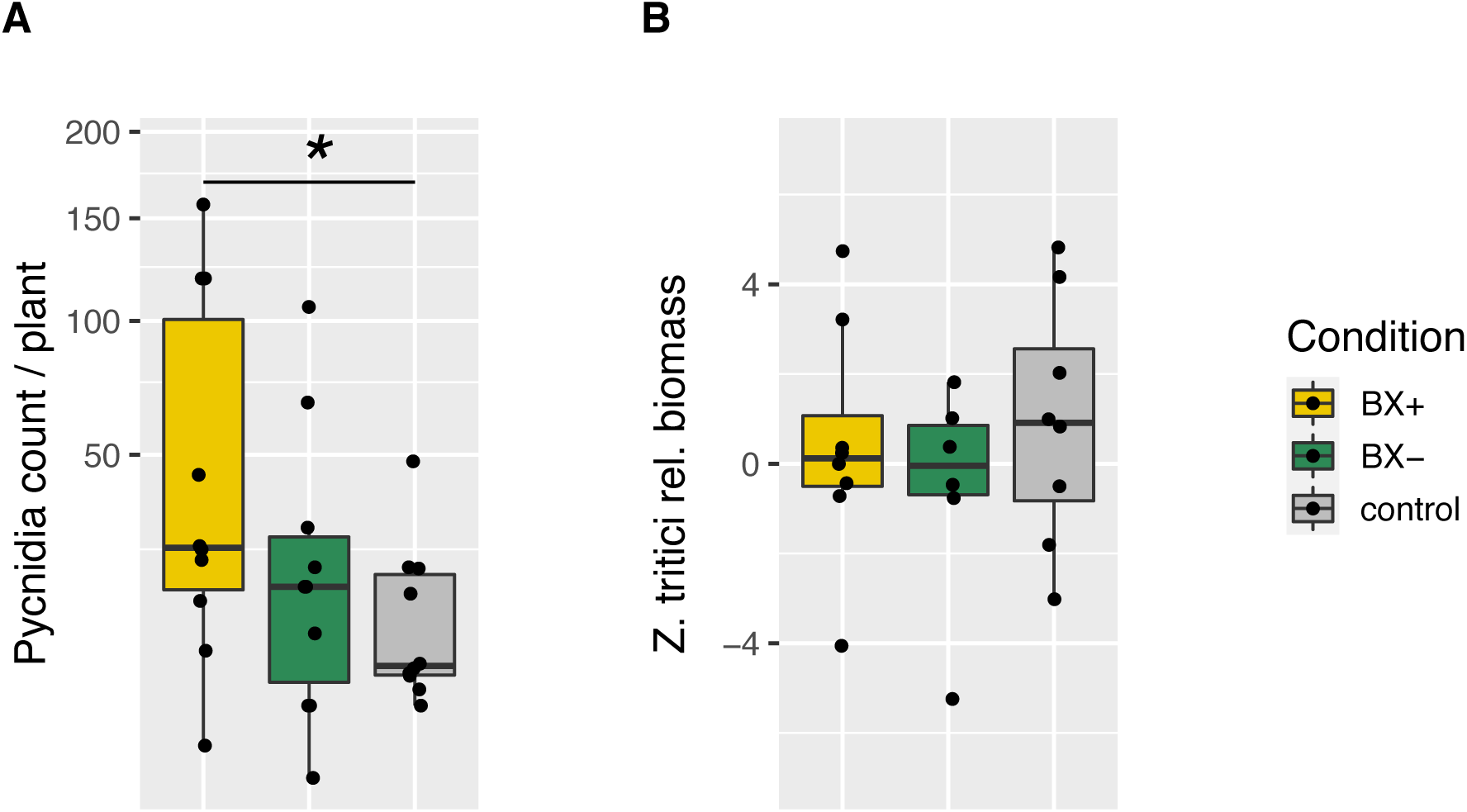
BX-effect on wheat pathogen resistance. Wheat plants from cultivar Drifter were grown on soils that were X-ray sterilized and re-inoculated with initial BX-conditioned bacterial microbiomes (‘BX+’ and ‘BX-’ soil variants), and on mixture of x-ray sterilized BX+ and BX- soil (‘control’ soil variants). Success of pathogen *Zymoseptoria tritici* colonization was measured as (A) pycnidia counts (mean per pot), counted on the infected leaf and (B) fungal biomass quantification by qPCR in infected leaf, calculated as relative to the mean of control plants. Data S5 documents the statistical analyses in detail. Stars reported on the graph correspond to significant difference with glm on square root data for (A) (significance code: *P* < 0.001 ***; *P* < 0.01 **, *P* < 0.05 *).

## Supplementary Tables

**Table S1.**
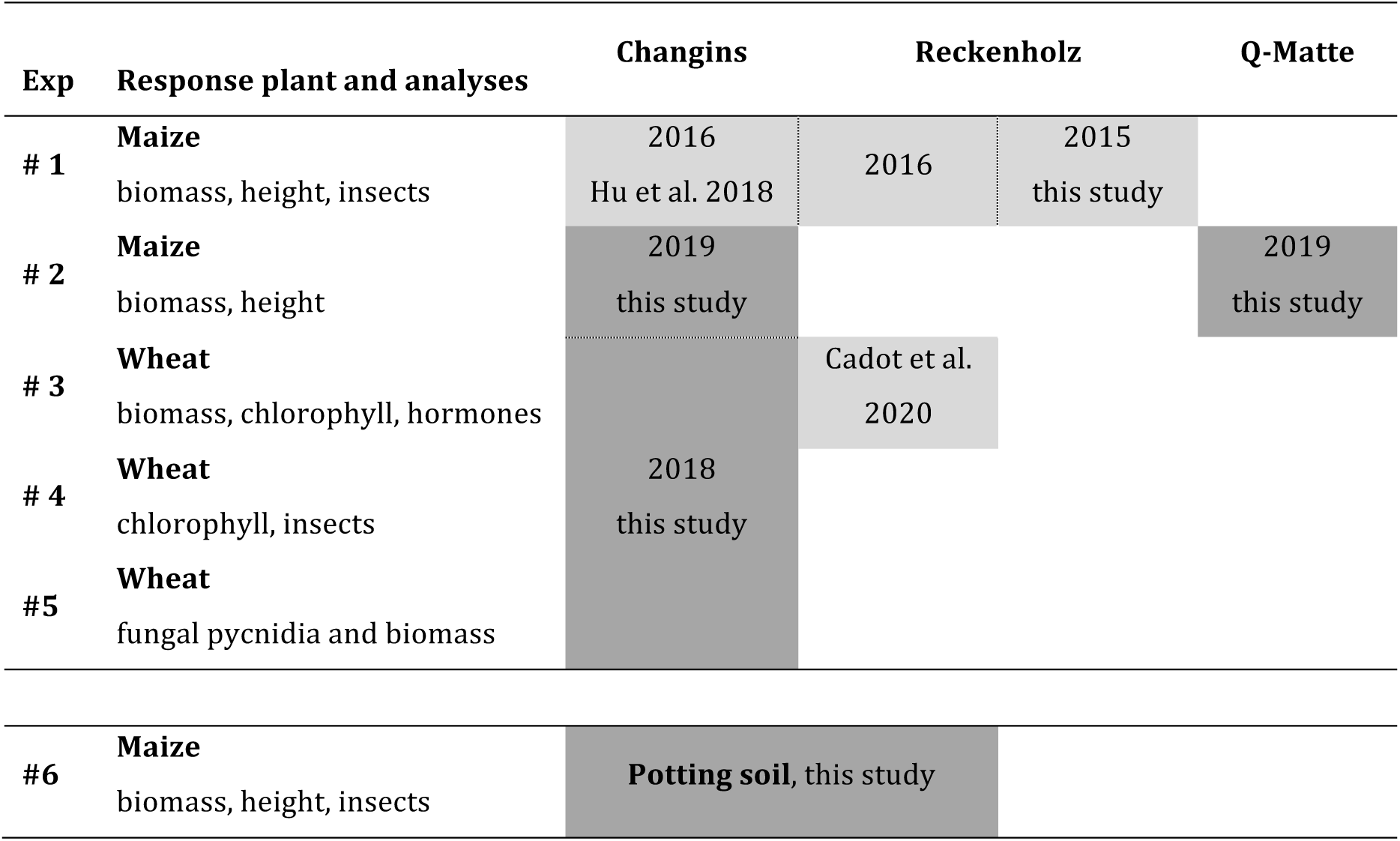
Experiments, analyses and batches of conditioned soils. Each feedback experiment (Exp) is listed with the response plant species and the performed analyses. The table details the different batches of conditioned soils (indicated as boxes) and their use in the different experiments. It also lists for each soil batch, the year of harvest, the type of conditioning (field = light grey; greenhouse = dark grey) and the reference, where the respective conditioning experiment was described. Of note, Experiment 6 was performed with potting and not field soil.

**Table S2.** Soil chemical characteristics. Physical and chemical characteristics of the three soils from the locations Changins, Q-Matte (Bern) and Reckenholz were determined in 1:10 water (‘…- H_2_O’, proxy for plant available nutrients) and 1:10 acetate-ammonium EDTA (‘…- EDTA’, proxy for reserve nutrients) extracts. Total iron was measured in nitric acid (‘Fe-HNO_3_’) extracts and soil texture was measured by fractionation. Letters indicate significant differences between locations (*P*-value < 0.05, Tukey HSD

|  |  | Changins |  |  | Q-Matte |  |  | Reckenholz |  |  |
| --- | --- | --- | --- | --- | --- | --- | --- | --- | --- | --- |
|  |  | mean | SD |  | mean | SD |  | mean | SD |  |
| P-H<sub>2</sub>O |  | 3.2 | 1.1 | b | 3.4 | 0.9 | ab | 5.3 | 0.4 | a |
| K-H<sub>2</sub>O |  | 44.8 | 5.0 | a | 24.1 | 0.3 | b | 23.0 | 1.6 | b |
| Mg-H<sub>2</sub>O |  | 14.0 | 0.4 | c | 15.8 | 0.5 | b | 20.5 | 0.3 | a |
| Ca-EDTA |  | 346.7 | 17.9 | a | 259.5 | 12 | b | 101.1 | 6.7 | c |
| NO<sub>3</sub>-H<sub>2</sub>O |  | 145.6 | 9.2 | a | 50.9 | 4.0 | b | 19.6 | 1.7 | c |
| NH<sub>4</sub>-H<sub>2</sub>O |  | 9.2 | 1.6 | a | 11.9 | 1.1 | a | 10.6 | 0.4 | a |
| Na-H<sub>2</sub>O | mg kg<sup>-1</sup> | 8.5 | 0.6 | a | 7.3 | 0.3 | a | 4.5 | 3.2 | a |
| Fe-H<sub>2</sub>O |  | 2.6 | 0.3 | b | 10.3 | 0.9 | a | 9.9 | 1.3 | a |
| Bor-H<sub>2</sub>O |  | 0.13 | 0.01 | b | 0.07 | 0.00 | c | 0.16 | 0.01 | a |
| P-EDTA |  | 64.6 | 4.4 | a | 25.4 | 1.8 | b | 63.2 | 2.8 | a |
| K-EDTA |  | 210.9 | 11.0 | a | 134.8 | 5.0 | c | 185.5 | 1.8 | b |
| Mg-EDTA |  | 177.0 | 11.7 | b | 189.1 | 7.5 | b | 513.0 | 14.2 | a |
| Ca-EDTA |  | 7883.7 | 622.5 | b | 9583.7 | 550 | a | 4601.0 | 153.0 | c |
| Mn-EDTA |  | 235.3 | 7.0 | b | 179.3 | 12.1 | c | 262.9 | 7.4 | a |
| Fe-HNO<sub>3</sub> | g kg<sup>-1</sup> | 23.2 | 1.1 | a | 23.3 | 1.2 | a | 20.9 | 1.0 | b |
| sand | % | 30.5 | 4.3 | b | 36.7 | 3.2 | ab | 37.5 | 2.4 | a |
| silt | % | 37.1 | 2.1 | a | 53.1 | 2.5 | b | 37.8 | 1.6 | a |
| clay | % | 30.5 | 3.4 | a | 10.2 | 0.8 | b | 24.8 | 0.8 | c |
| pH |  | 7.8 | 0.4 | a | 6.6 | 0.4 | b | 6.9 | 0.4 | b |

